# Tunable DNA Origami Nanosensors for Detection of Multiscale Spatial Ion Concentration Gradients

**DOI:** 10.1101/2025.10.16.682652

**Authors:** P Beshay, Z Osborn-King, M Kruse, J Marshall, T Teng, J Song, C Castro, BA Walter

**Affiliations:** The Ohio State University, Department of Mechanical and Aerospace Engineering; The Ohio State University, Department of Biomedical Engineering; Ohio State University; The Ohio State University

**Keywords:** DNA Nanotechnology, Ion sensing, Concentration Gradients, Microfluidics, FRET

## Abstract

Ion gradients play a vital role in cellular signaling, mechanobiology, and organ-level homeostasis. Despite their importance, accurately mapping these spatial gradients at biologically relevant length scales remains a challenge due to the limited tunability and spatial resolution of conventional fluorescent sensors. Here, we present a DNA origami-based sensor (NanoDyn) with tunable sensitivity that enables the detection of Na^+^ ion gradients across micron to millimeter scales. The sensor design leverages programmable DNA base-pairing interactions to control both the detection range and sensitivity of the sensor. Using fluorescence spectroscopy, we show that NanoDyn can exhibit programmable sensing ranges spanning ∼100 – 1675 mM Na^+^. To validate the ability to quantify ion gradients and investigate its spatial resolution, we use a custom microfluidic gradient generator, showing that NanoDyn can resolve changes in ion gradients across multiple scales and over distances as little as ∼6 µm, which here, is limited by the resolution of the microfluidic device. In highlighting the potential of DNA nanodevices as multiscale tunable ion gradient sensors, together with their biocompatibility, high temporal resolution, and potential for multiplexed functionalization, this work expands on the role that DNA nanodevices can play in spatial sensing to study ion-mediated processes in microenvironments. Overall, this work advances DNA nanotechnology as a versatile foundation for biosensing with capabilities to probe ion-mediated signaling in health and disease.

## Introduction

Ion gradients are fundamental regulators of biological processes across multiple scales, with micron- to millimeter-scale gradients playing key roles in the normal function and homeostasis of cells [1-3], tissues [4-6], and organs [7, 8]. For example, the kidney relies on precisely maintained sodium gradients for its essential functions like filtering waste products or regulating electrolyte levels in the blood [9, 10], and in load bearing musculoskeletal tissues, including the intervertebral disc (IVD) and articular cartilage, ionic gradients develop from compression of charged and hydrated tissue, which in turn regulates tissue hydration, mechanical response, and mechanobiology down to the cellular level [11-13]. The fundamental role of ion gradients in regulating biological processes across multiple scales underscores a need for advanced measurement systems that provide high sensitivity and spatiotemporal resolution, enabling real-time monitoring of the ionic microenvironment at micron to millimeter scales.

Advancements in molecular biosensors have enabled novel techniques for measuring ion concentrations in biological systems. Some common techniques include fluorescent indicators where an ion-binding site is attached to a fluorophore molecule or electromechanical sensing platforms, such as micro ion flux estimation (MIFE) and electrical sensors [14-17]. Fluorescent indicators have the advantage of being highly sensitive and easy to implement, but have limited reversibility and tunability, and their narrow fluorescence spectra and spectral overlap with other sensors can limit some applications [14, 18, 19]. Additionally, the high specificity of dye to certain ions can be a major strength for certain applications, (e.g., Ca2+) but a challenge if interested in other ionic species where there are less available options [20]. Electromechanical techniques can facilitate localized and real-time detection of ionic activity in biological microenvironments [21-23], yet are subject to other technical challenges associated with limited spatiotemporal resolution and complex calibration. Collectively, both approaches are impeded by factors such as invasiveness, limited tunability and/or spatiotemporal resolution, and calibration complexity, restricting their application for high-resolution spatial gradient mapping across multiple scales in complex biological systems. These factors highlight a need for a sensor that can overcome some of these limitations.

Recent advances in DNA nanotechnology have opened new avenues for the design of dynamic devices that are ∼1-100 nm in size with tunable response to a variety of inputs [24, 25]. In particular, the DNA origami method [26] enables the self-assembly of 2D and 3D nanostructures with diverse geometries and programmed dynamic behaviors [25, 27]. The ability to control DNA interactions in a sequence specific manner has allowed the functionalization of DNA devices with various molecules, such as proteins, peptides, lipids, aptamers, and small molecules with nanometer precision [28-30]. This, along with their biocompatibility, has enabled the use of DNA-based devices in a host of biological applications, from drug and gene delivery to imaging and biosensing [31, 32]. As biosensors, DNA-based devices have been used for detecting a variety of signals such as nucleic acids [33], proteins [34], small molecules [35], and changes in pH [36] with a high sensitivity [37]. In addition, prior work has demonstrated DNA devices that are sensitive to ion concentrations [38-42]. However, these have limited tunability since they only rely on sequence approaches to tune sensitivity (e.g. quadruplex or i-motif). Furthermore, these prior studies with DNA-based sensors have explored detection either in bulk (i.e. over an entire solution), at single molecule level, or in cellular compartments. The ability to tune sensitivity over a large range required for sensing at multiple scales and the mapping of spatial concentration gradients remain a challenge.

Here, we present a DNA-based sensing platform that is capable of measuring Na+ ion gradients with widely tunable sensitivity and versatile spatial resolution spanning microns to millimeters. We leverage structural and sequence design parameters that control the conformation state and the flexibility of the device to tune the sensitivity and detection ranges of the sensor. To this extent, we designed a microfluidic device to generate ion gradients with high precision. We utilize fluorescence spectroscopy to show that the sensors of different parametric designs have effective ranges that span from ∼100-238 mM on the lower end, to ∼100-1675 mM on the higher end. We also leverage our microfluidic device design and confocal microscopy to show that the sensors are capable of measuring changes in Na+ gradients as small as 25 mM at the micron scale. Overall, this sensing platform offers both high spatial resolution and tunable sensitivity characteristics, providing new tools that could help in understanding ion-mediated biological processes.

## RESULTS & DISCUSSION

### NanoDyn Device Design and Parameter for Tuning Ion Sensitivity

This study implements a previously reported dynamic DNA origami device referred to as the NanoDyn [43, 44], which consists of two barrel components, each ∼50 nm long, connected by six long linkers joining the two barrels together (Figure 1A). Of the six linkers, 5 are classified as modulating linkers (ML), and are folded as either single-stranded (SS) DNA or are folded to have two duplex regions so the linker can flexibly extend (i.e. unconstrained (UNC) linker); duplex regions of the UNC linkers are generally more rigid than the SS linkers and allow more defined relative motion of the two barrels. The sixth linker is classified as a fluctuating linker (FL) which can bind with itself to form a closed loop holding the two barrels together. The FL also has a Cy3/Cy5 Förster Resonance Energy Transfer (FRET) pair for readout of this binding. The strength of the DNA binding interaction of the FL can be modified by varying base-pair (bp) lengths (*N*_*bp*_). The device has two primary stable states, an open state where the two barrels are loosely tethered together by the linkers, and a closed state where the two barrels are held together by the DNA binding interaction of the FL. The closed or open state of the device can be determined by FRET efficiency of the FL.

**Figure 1.**
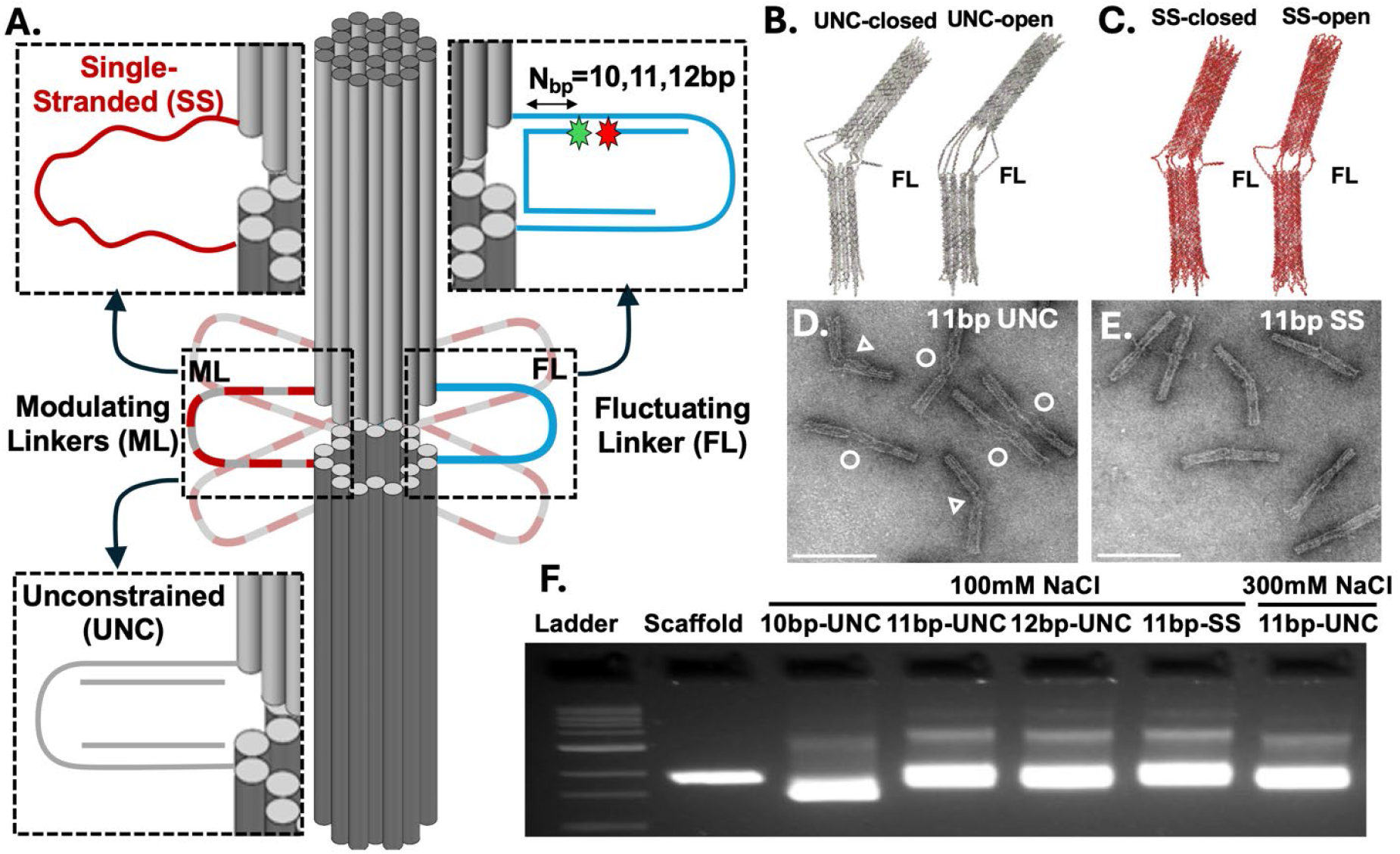
Sensor Design Variations. **A**) Schematic of the NanoDyn structure. Five modulating linkers (ML) and one fluctuating linker (FL) connect the top and bottom barrels. The modulating linkers are either unconstrained (UNC) or single stranded (SS) depending on the version. The number of base pair interactions (N_bp_) of the fluctuating linker can vary and for designs used here ranges from 10 to 12 base pairs. (**B**,**C**) Average configurations from coarse grained molecular dynamics simulations of NanoDyn devices in both closed and open states for UNC and SS designs. (**D**,**E**) TEM images of the 11bp-UNC and 11bp-SS; Scale bar = 100nm. 11bp-UNC closed structures are marked with a triangle (Δ) and open structures are marked with a circle (O), and it is not possible to differentiate open and closed structures for 11bp-SS structures. **(F)** Gel electrophoresis confirms structures are well folded and stable at lowest testing concentration of 100mM NaCl with 1mM MgCl_2_ for one hour.

Ion concentration is an important factor influencing the stability of DNA base-pairing interactions [45]; hence, the stability of the closing interaction should depend on the local ion concentration in solution. This work focuses on engineering the sensitivity of device conformations to local ion concentration to enable measurement of cation, specifically Na+, concentration gradients. To optimize the NanoDyn sensor for a physiologically relevant range of NaCl concentrations, two design parameters were varied. First, *N*_*bp*_ was varied, which mediates the stability of the closed state. Second the other 5 modulating linkers (ML) were folded in either an UNC or SS state, which modulates the flexibility of the NanoDyn especially in the open state. To visualize the effect of the ML, we simulated two versions of the NanoDyn: one with all 5 MLs in the SS state and another with all 5 MLs in the UNC state, in both closed and open configurations using the coarse-grained molecular dynamics model oxDNA [46, 47]. Figures 1B-C show average configurations from these simulations, revealing a large difference in barrel separation between the open and closed states for the UNC case, consistent with the expected flexibility. In contrast, for the SS case the separation between the barrels is similar for the open and closed states, likely due to the entropic elasticity of the SS linkers. This suggests the NanoDyn with SS linkers would likely transition back to the closed state more readily, leading to higher FRET values on average.

Our prior work with NanoDyn devices suggests devices with *N*_*bp*_ = 13 bp are stably closed [44], while devices with *N*_*bp*_ = 10 bp are largely open [43]. Here we aimed for a dynamic device that fluctuates between open and closed states where the NaCl concentration would affect the stability of the closed state. Hence, we decided to test *N*_*bp*_ = 10, 11, and 12 bp interactions on the FL, and we chose the middle interaction length, *N*_*bp*_ = 11, to test the effect of the MLs being folded in the UC or SS state. In total, four variations of the device were tested, which we denote in terms of the *N*_*bp*_ and ML configuration: 10bp-UNC, 11bp-UNC, 12bp-UNC, and 11bp-SS. All four NanoDyn versions were folded, purified, and characterized by transmission electron microscopy (TEM) imaging (details in Methods). TEM images of all the versions (Supplemental Figure 1) confirmed well-folded nanodevices exhibiting a distribution of configurations. The 11bp-UNC devices (Figure 1D) in particular, showed a distribution of open and closed states that were consistent with simulations and distinguishable based on the relatively larger gap between barrels for the open state, while closed devices exhibited closer barrels with a kink at the FL. In contrast for the 11bp-SS devices, we did not observe any clearly open structures in TEM images (Figure 1E). This could either mean they are all closed, or that it is not possible to distinguish open and closed states, which is consistent with simulation results suggesting both open and closed SS structures have a small gap between the barrels. Finally, based on the intention to use these devices to measure a range of NaCl concentrations, we tested the stability of these designs at the lowest NaCl concentration used in this study (100 mM NaCl). Agarose gel electrophoresis (AGE) results (Figure 1F) show all devices exhibited a sharp clear band that ran similar to or faster than the scaffold, indicative of intact DNA origami structures. The 10bp-UNC ran faster, which is also consistent with prior work showing more open NanoDyn devices run faster in AGE experiments than more closed devices [43]. These results confirm that devices will remain stable in experimental cation conditions.

### Tuning Cation Concentration Sensitivity of NanoDyn Sensors

The Cy3 donor and Cy5 acceptor on the FL provide a quantifiable readout of the closed/open state of the NanoDyn, with low FRET indicating an open state (i.e. donor and acceptor far apart, Figure 2A top) and high FRET indicating a closed state (i.e. donor and acceptor close together, Figure 2A bottom). Figure 2A shows schematics and TEM images for both open and closed NanoDyn devices (TEM images show 11bp-UNC structures). The FRET response for all four variations of the device was measured at 9 different salt concentrations over the range of 100-2000 mM NaCl, which contains physiologically relevant range between 100-350mM NaCl [48-50]. Figure 2B compares the FRET response of three versions of the NanoDyn, all with 5 UNC linkers while *N*_*bp*_ is varied to be 10, 11, or 12 bp (i.e. 10bp-UNC, 11bp-UNC, and 12bp-UNC versions) and Figure 2C compares the FRET response for *N*_*bp*_ = 11 bp while varying the ML linkers to be either SS or UNC (i.e. 11bp-UNC and 11bp-SS versions). For all versions there is a clear increase in FRET with increasing NaCl concentrations, with higher FRET values observed for higher *N*_*bp*_ and for the SS configuration of the ML linkers. The FRET readout for the 11bp-UNC and 12bp-UNC versions saturates around an NaCl concentration of 600 mM, while 10bp-UNC approaches saturation at 2000 mM. The 11 bp-SS design exhibits the highest FRET values and saturates earlier around a concentration of 300 mM (Figure 2C), which is consistent with the simulation result that suggested SS designs would likely transition back to the closed state more readily. We applied exponential curve fits to the FRET response curves to determine an NaCl exponential concentration constant, or saturation constant *C*_*sat*_, and an initial sensitivity (i.e. slope of curve fit at an NaCl concentration of 100 mM), which are reported in Figures 2C-D, Supplemental Table 3. We estimated an effective measurement range as two times the *C*_*sat*_, which yielded high end measurement ranges of 1675mM, 365mM, 448mM, and 238mM for the 10bp-UNC, 11bp-UNC, 12bp-UNC, and 11bp-SS, respectively (shown as shaded region in Figures 2B-C, Supplemental Table 3). These results indicate that the 10bp-UNC provides a low sensitivity and large measurement range, the 11bp-UNC and 12bp-UNC provide intermediate sensitivity and intermediate measurement range, and the 11bp-SS provides a high sensitivity with low measurement range, highlighting the ability to tune sensing parameters for desired applications. In particular, we selected the 11bp-UNC and the 11bp-SS for subsequent experiments focused on testing capabilities to measure cation concentration gradients across biologically relevant length and concentration scales.

**Figure 2.**
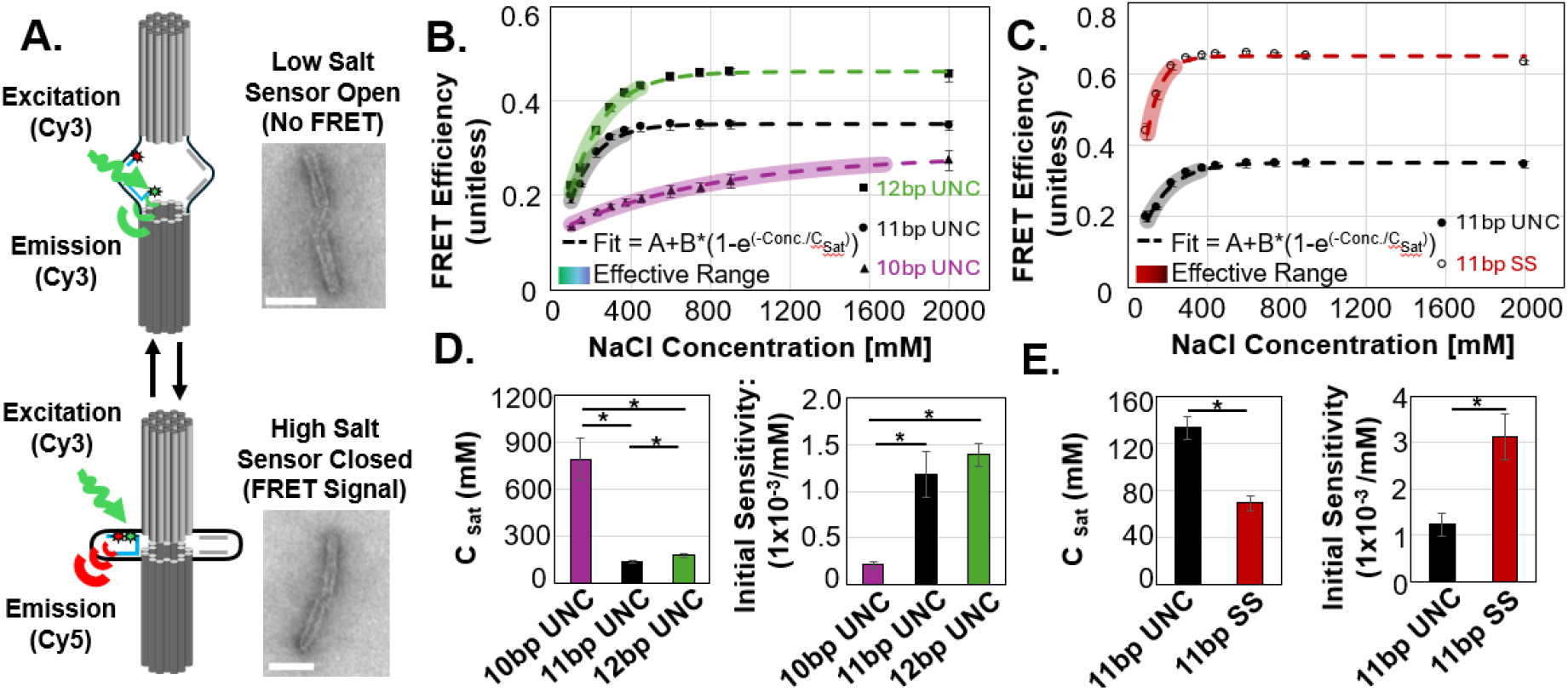
Tuning Ion Concentration Sensitivity of NanoDyn Sensors. (**A**) Schematic of the conformational change of the NanoDyn in response to salt concentrations. In low salt concentrations sensors are open (top), resulting in the Cy3 and Cy5 fluorophores being separated and a lower FRET efficiency. In high salt concentration the sensors are closed, resulting in the Cy3 and Cy5 fluorophores being closer together and a higher FRET efficiency. TEM images show examples of open and closed 11bp-UNC structures. Scale bar is 50 nm. (**B, C**) Demonstration of how the design parameters (bp or UNC/SS) impact the concentration dependent changes in FRET efficiency. Data are fit with the equation FRET=A+B*(1-exp(-Concentration/C_sat_), where A, B and C_sat_ are fitting constants. Shaded regions represent the ‘effective range’ of each sensor. (**D, E**) Quantification of how design parameters (bp or UNC/SS) impact the saturation constant (C_sat_), i.e., how quickly the FRET signal saturates with increasing concentration, and the initial sensitivity. Initial sensitivity was determined by evaluating the derivative (i.e., d(FRET)/dConc) at c=100mM. *=p<0.05.

### Multiscale Spatial Sensitivity of NanoDyn Devices

#### Macroscale

The response of NanoDyn sensors to spatial changes in NaCl concentration over the mm-scale was investigated using a 2-inlet microfluidic gradient generator which mixes laminar flows and then recombines them into a single chamber with 6 mixed inlet flows (Figure 3A) [51]. NanoDyn sensors (11bp-SS or -UNC) at a concentration of 25nM were combined with either 150mM or 200mM NaCl and flowed into the microfluidic device at a flow rate of 1uL/min. TEM imaging demonstrated that Nanodyn sensors remained intact after flowing through the microfluidic device (Supplemental Figure 4). The FRET efficiency of the NanoDyn sensors within each of the 6 inlet flow regions was assessed and correlated against either distance across the microfluidic chamber or the interpolated NaCl concentration (Figure 3B). The slope of the linear relationship between FRET Efficiency and either distance or interpolated concentration, representing both spatial detection and concentration sensitivity, was compared between sensor designs. Briefly, the spatial variation in interpolated NaCl concentration was calculated using a scaling factor determined by flowing red-dye and water alone (no red-dye) into the microfluidic gradient generator. The inlet red-dye concentration was chosen so that the color intensity scaled linearly to concentration via Beer-Lambert’s law, and the red-dye intensity across the gradient chamber was normalized from zero to one. The normalized dye intensity was scaled to the NaCl inlet concentrations (0 corresponding to 150mM NaCl and 1 corresponding to 200 mM NaCl) to generate the interpolated NaCl Concentration. Clear differences in FRET efficiency were observed across the chamber for both sensor designs (Figure 3A, right) and FRET efficiency linearly correlated with both distance across the channel and with the interpolated NaCl concentration (Figure 3B). The slope of the FRET v. distance and FRET v. interpolated concentration correlations was at least twice as large for the 11bp-SS sensor than the 11bp-UNC sensors indicating that 11bp-SS has a higher sensitivity to lower NaCl concentrations than 11bp-UNC. This aligns with prior observations from fluorometer experiments and suggests a higher spatial sensitivity to cation concentrations within smaller concentration gradients.

**Figure 3.**
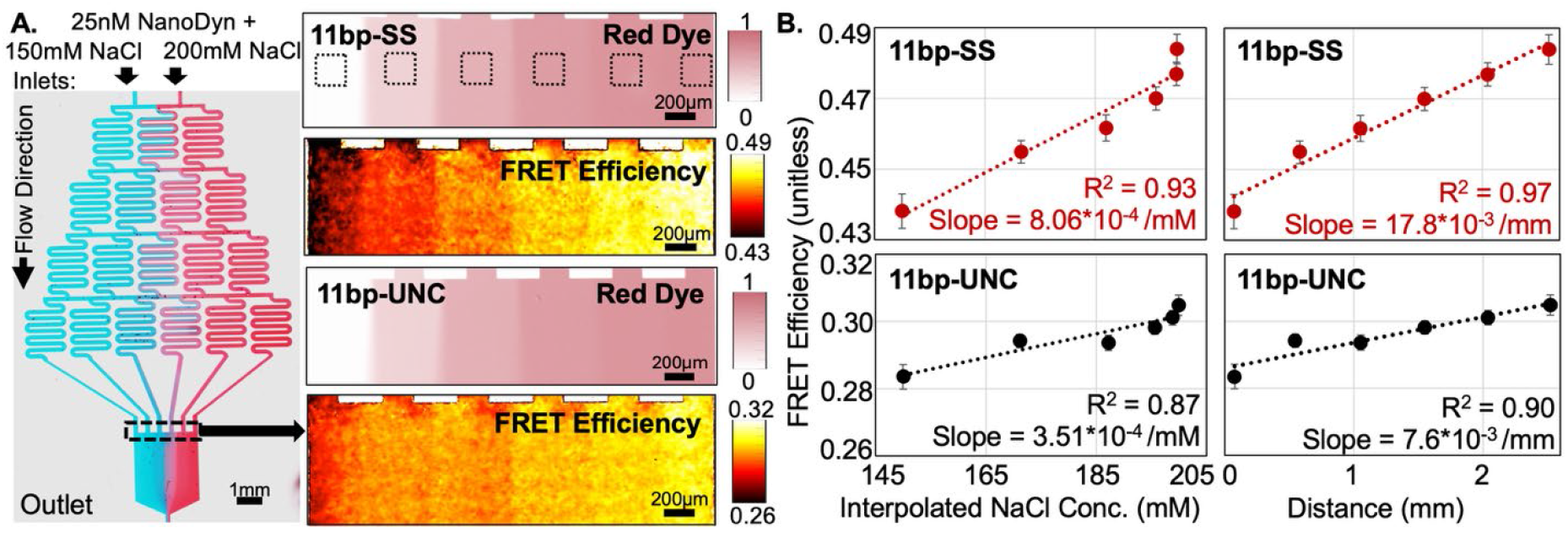
Macroscale Spatial Resolution of NanoDyn Sensors. (**A**) (**Left**) Representative image of the 2-inlet device used for these studies using red and blue colored dyes, with the corresponding imaged region shown with the black, dotted rectangle. Salt concentrations of 150mM on the left inlet and 200mM on the right inlet are used for all the following experiments. **(Right)** Stitched Images of the red dye and FRET image taken within the same device. 186 x 186um (150 x 150 pixel) areas are shown as dashed line boxes in the top image to visualize the areas used for averaging in 3B. **B**. Plots correlating each sensor type to the interpolated sodium gradient across the distance of the channel. 11bp-SS sensors have a higher slope in both categories, due to their stronger concentration sensitivity.

#### Microscale

To determine the spatial sensitivity of NanoDyn sensors to NaCl concentrations at length-scales relevant for cellular applications, the sensor response was evaluated in a microfluidic device where two laminar flows converge without mixing, allowing the spatial transition between two known concentrations to occur over micron-scale distances. Four concentration gradients, increasing from 150mM by 300mM, 200mM, 50mM, and 25mM, were used for both 11bp-SS and 11bp-UNC sensors. An area encompassing the transition between flows was analyzed (186 x 186 µm, Figure 4A). To characterize the spatial sensitivity, the mean peak slope of the transition was calculated for each replicate trace and averaged (Figure 4B, on the plots and Supplemental Table 4). In addition, a distance to equilibrium was calculated for each sensor/concentration combination to serve as an approximation of the minimum spatial resolution. This equilibrium distance was calculated by dividing the average change in FRET between the two known concentrations by the mean peak slope. Results demonstrate that at a smaller change in concentration (i.e., Δ25mM or Δ50mM) the 11bp-SS had greater spatial sensitivity (i.e., slope) than the UNC sensors (Figure 4B) and were able to differentiate 25mM over 8-10µm. At greater concentration changes (i.e., Δ200mM or Δ300mM) the UNC sensors had greater sensitivity and were able to detect a change of 300mM over ∼6µm. When assessing the spatial range at which all sensors were able to detect concentration changes, the range was ∼6.5-13µm, which suggests the sensors are suitable for measuring cation concentration gradients at cellular length scales (Figure 4B and Supplemental Table 4).

**Figure 4.**
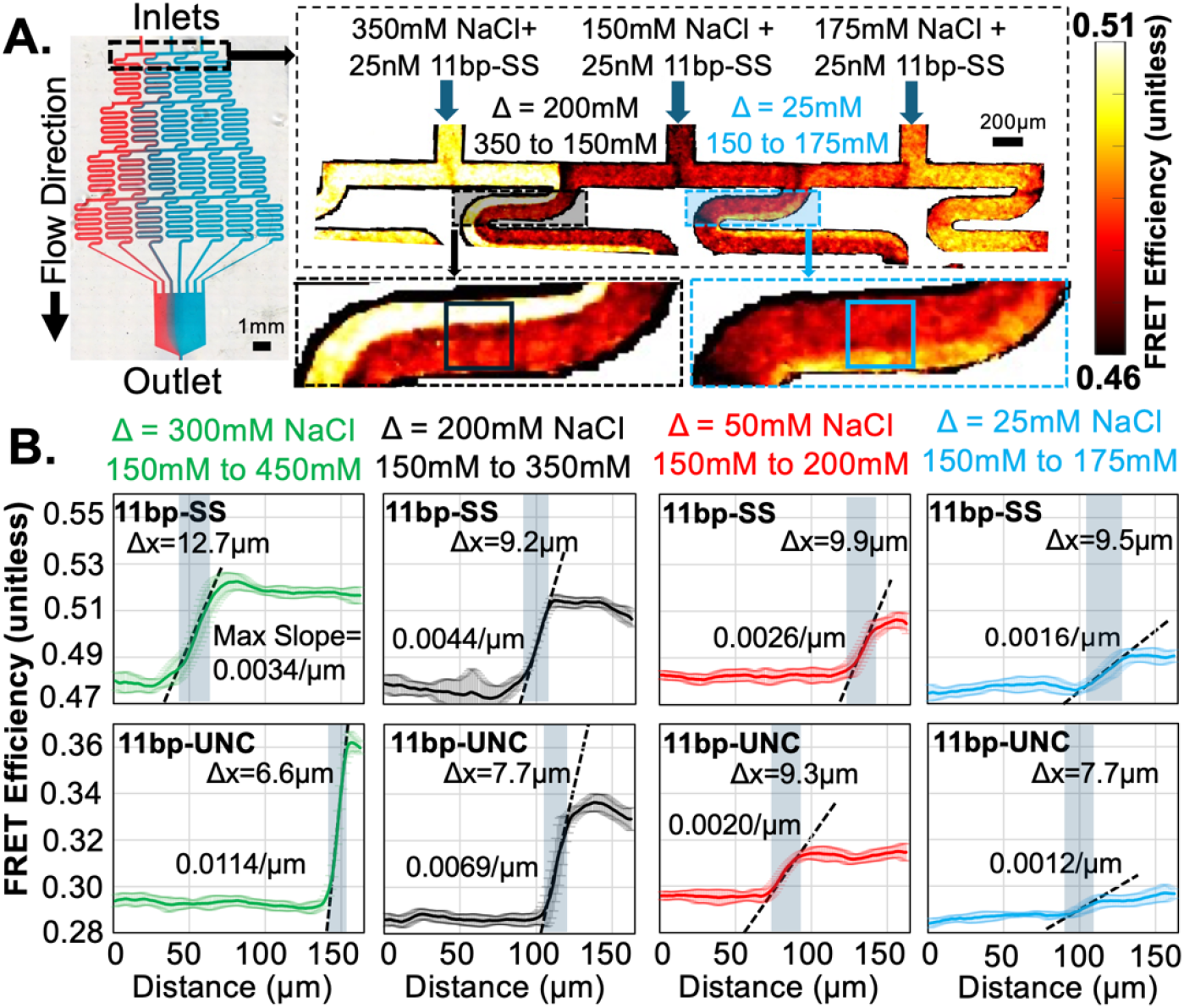
Microscale Spatial Resolution of DNA Origami (DO) Sensors. (**A**) (Left) Representative image of microfluidic device with three inlets used to establish abrupt changes in NaCl concentrations. Zoom in shows the FRET image illustrating visual changes in FRET and highlighting boxes where quantification was performed. (**B**) Quantification of the spatial resolution of 11bpSS & 11bpUNC DO across laminar flows of varying NaCl concentrations within a microfluidic device. The channel width is 200µm. Dashed line in B. is the max slope of the transition between two adjacent laminar flows and shaded region is the transition distance between two concentrations. Results demonstrate DO have spatial sensitivities that can detect changes in cation concentration of 25mM or more at the micron/cellular scale (∼6.5-13µm).

## CONCLUSIONS

In this study, we presented a DNA origami sensing platform that enables tunable ion sensing across a wide region of Na^+^ concentrations, spanning from ∼100 – 1500 mM. By systematically changing the folding configurations and base pairing interactions on the linkers of NanoDyn, we established a structural design strategy to tune both the sensitivity and the detection range of the sensors. We used a microfluidic gradient generator to show the spatial sensitivity of NanoDyn to cation concentration at both the macro- and microscales. Our findings show that NanoDyn sensors can sense changes in Na^+^ as little as 25mM across millimeter distances and with a spatial resolution as little as ∼6 µm, demonstrating feasibility to assess physiologic changes in ion concentration on the micron scale. The spatial resolution measured here is limited by the microfluidic device, and it is possible the sensor could measure concentration gradients at smaller length scales, even down sub-micron as demonstrated in single molecule detection assays [52, 53]. Additionally, these microfluidic experiments demonstrate the capacity to apply DO sensors suspended in a flowing solution, expanding on prior applications that have ion sensitive DNA-based devices for fixed measurements in space or in bulk (i.e. over an entire solution) [38, 39, 41, 54]. This also circumvents challenges like photobleaching since sensors are constantly renewed in the area of interest. In general, the application of sensors within a flow could be synergistic with approaches where individual sensors are immobilized on uniform or patterned surfaces [55, 56].

Recent studies on DNA-based structures show a response time to ion changes on the scale of milliseconds and that these changes are reversible [42], suggesting these nanosensors could also be used to assess dynamic changes in ion gradients over time with fast time resolution. However, future studies are required to explore important factors like long-term stability and reversibility in complex environments. Regardless, their biocompatibility and high programmability make these nanosensors excellent candidates for integration in complex biological systems [57], on cell surfaces [58, 59], or in tissue microenvironments [60] for example. Overall, our work underscores the potential of dynamic DNA nanodevices to be deployed as tunable nanosensors to address challenging questions regarding the role of ion gradients in complex micro-scale environments. Looking forward, such devices can serve not only as measurement tools to study disorders related to disruptions in ion concentrations, fluxes, or gradients, but also as research probes to investigate ion-mediated processes across multiple length scales [3, 6, 8, 61].

## EXPERIMENTAL METHOD

### Device Fabrication, Purification, and Simulation

The NanoDyn structure has been previously reported [43, 44] and the structure design has been shared on an online design database (https://nanobase.org/structures/19) [62]. The structure was folded using an M13mp18 derived 8,064 nucleotide scaffold [63] prepared in house as previously described [64], and folding was carried out similar to previously established methods [64, 65]. For folding, 150uL of 100 nM scaffold DNA was combined with 300uL of DNA staple strands, each at a concentration of 500 nM, 75uL of 10x folding buffer (FOB; 10 mM of EDTA (ethylenedinitrilo)tetraacetic acid), 50 mM Tris (tris(hydroxymethyl)aminomethane) (Millipore Sigma CAS-77861) and 50mM NaCl) was added to 75uL of 180 mM MgCl_2_ (Millipore Sigma CAS-7786303) and 150uL of ddH_2_0 for folding in a total reaction volume of 750uL. The staples added depended on the version of the structure that was being folded. The staples used to form the two barrels of the structure (Supplemental Table 2) were the same across all versions while the staples incorporated to form the fluctuating and 5 modulating linkers varied across different version of the device (Supplemental Table 1). Final concentrations of the folding reaction include 20 nM scaffold, 200 nM staple strands, 1 mM EDTA, 5 mM Tris, 5 mM NaCl, and 18 mM MgCl_2_. The folding reactions were then placed in a temperature-controlled water bath at 70°C for 30 minutes for a melting phase before rapidly switching them to a separate temperature-controlled water bath at 52°C for an annealing phase for 1 hour. Once folded, structures were stored in 4°C until use.

Proper folding was first confirmed by agarose gel electrophoresis as previously described [64] using a gel consisting of 2% agarose in 0.5x TBE buffer (Tris, borate, and EDTA) buffer containing 45 mM boric acid, 45 mM Tris, 1 mM EDTA with 11 mM MgCl_2_, and 0.5 μg/mL ethidium bromide. After structures were verified as well-folded, the structures were purified by centrifugation in the presence of polyethylene glycol (PEG), similar to prior work [66]. Briefly, the structures were mixed in equal volumes at a 1:1 ratio of sample to a solution with 15% PEG 8000 (w/v) in 500 mM NaCl (for a final PEG concentration of 7.5%) before centrifugation at 16,000g for 35 min. The supernatant was removed and the pellet was collected and resuspended in phosphate buffered saline solution with either 200 or 300 mM NaCl with 1mM MgCl_2_. This procedure was repeated a second time to ensure effective removal of excess staple strands. The 200 mM NaCl concentration for resuspension was used for experiments in 100 mM NaCl (after a 2-fold dilution), and the 300 mM NaCl concentration was used for all other experiments.

CaDNAno design files were converted to oxDNA configuration and topology files for simulation using tacoxDNA [67] built in oxView [68]. The simulation process includes relaxation and normal simulation. After relaxation with applying mutual traps, the oxDNA2 interaction model was used to conduct coarse-grained simulations. A total of 10^8 steps with GPU acceleration were used and each step was set to 15.15 fs. Simulation parameters included an Anderson-like thermostat, temperature at 30 °C, and monovalent salt concentration at 0.5 M, all standard conditions in oxDNA simulations. The processes mentioned above were executed through a shell script for all structures in this study in a Linux computer equipped with a NVIDIA GeForce 3090 graphics card. The trajectory file was later analyzed in oxDNA analysis tools written by Python, including mean configurations, root-mean-square deviation (RMSD), and root-mean-squared fluctuations (RMSF). The mean configurations were then generated in oxView.

### TEM Imaging

TEM Imaging samples were prepared following previously established protocols. Copper mesh grids (400 mesh, Ted Pella’s Formvar/Carbon grids (01754-F) were plasma-treated using a PELCO easiGlow Glow Discharge Cleaning System. A drop of purified sample (12 μL) was then deposited on the grid and incubated for 6 min. Excess sample was wicked away with filter paper. Samples were negatively stained with uranyl formate (UFo) (2%) by adding a droplet of 10 μL to parafilm, gently placing the grid in the droplet and immediately wicking the droplet away. This was followed by deposition of a second 20 μL UFo droplet that was incubated on the grid for 40s before removal using filter paper. The samples were then allowed to dry for at least 15 min prior to imaging. Images were collected with FEI Tecnai G2 Bio Twin TEM at 80 kV.

### Fluorescence Characterization of NaCl Sensitivity

The emission spectra were collected using a Horiba scientific FluoroMax-4. First, the donor (Cy3) was excited directly while the donor and acceptor (Cy5) emissions were measured. The excitation wavelength was centered at 510nm with a width of 5nm while the emission was measured between the range of 530nm-750nm with a step size of 5 nm. Second, the acceptor was excited directly while only measuring the emission of the acceptor. The excitation wavelength was centered at 600nm with a width of 5nm while the emission was measured over the range of 620nm-750nm with a step size of 1 nm. From the fluorescence emission spectra, the FRET efficiency [69] was calculated using a custom MATLAB code. To calculate FRET efficiency, the intensity of the acceptor excited by the donor (*I*_*A/D*_) was divided by the sum of the intensity of the donor excited directly (*I*_*D/D*_) and the intensity of the acceptor excited by the donor *I*_*A/D*_. All intensities were calculated over a 20nm range. Equation 1 below depicts the calculation used for FRET efficiency.

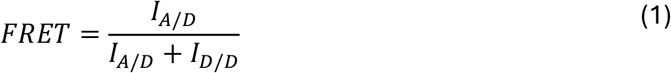

For all intensity measurements, a blank of the intensity at 300 mM NaCl was subtracted to account for noise in the measurement; 300 mM NaCl was chosen because it was the base buffer for all structures measured on the fluorometer.

### Microfluidic Device Fabrication and Design

PDMS (Sylgard) utilized for making microfluidic devices was mixed using a 10:1 ratio of elastomer base to curing agent. This mixture was then desiccated, poured into a petri dish containing a silicon wafer etched with the patterned microfluidic design (Micro Lithography Services, Essex, UK), and cured for 2 hours at 60°C. Cured PDMS was then removed from the petri dish and individual microfluidic devices isolated. For each device, inlets and outlets were punched using a 1mm biopsy punch (Integra Life Sciences) and then bound to #1 glass coverslips using a plasma bonder (Harrick Plasma) after clearing both PDMS and glass surfaces of dust particles using Scotch tape. To facilitate flow into the device, three segments of 19G needles (EXCEL, ID = 0.686mm/0.027in) were bent to form a ∼90o angle. These needles serve as the connection between the clear polymer tubing (Cole-Parmer, ID = 0.381mm/0.015in) and the microfluidic device. New tubing was cut with each experiment to ensure no contamination of reagents between experiments.

### Imaging DNA Origami Inside of Microfluidic Device

After device fabrication, each device was flushed of air bubbles using 2 syringe pumps (Harvard Apparatus) with plastic syringes connected to the microfluidic via tubing and needle connectors. The tubing and device were flushed with NaCl solution containing no sensors, with each inlet set at the NaCl concentrations intended for each experiment. 50nM 11bp-SS or 11bp-UNC sensors suspended in 300mM NaCl were heated at 37°C for ∼10 mins prior to use within the device to minimize aggregation, then mixed 1:1 with a NaCl solution to obtain 25nM of DNA origami sensors at the desired NaCl concentration following Equation 2:

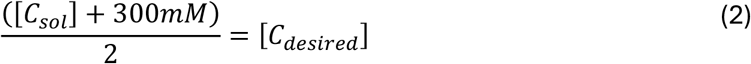

Where C_sol_ is the concentration of the NaCl solution mixed with the sensors and C_desired_ is the desired concentration for experimentation. After repeating this process for each remaining inlet of the microfluidic device, the 25nM sensor mixture was then loaded into 500uL glass syringes (Hamilton) and loaded into the device at a flow rate of 15uL/min for 9 minutes. After sensor loading, the device was allowed to equilibrate for 20 mins at 1uL/min to ensure a steady-state gradient profile before imaging. Sensors were imaged using a Nikon A1R microscope with the following settings: Channel 1) Cy3 excited with a 561 nm laser line emission captured with a bandpass filter (570-613 nm), Channel 2) Cy3 excited with a 561nm laser line and Cy5 emission captured with a bandpass filter (663-738 nm), and Channel 3) Cy5 excited with a 638 nm laser line and emission captured with a bandpass filter (663-738 nm). All images were acquired at 20% laser power, Galvano scanning, and using 16 lines for averaging. Gradient and Inlet regions were recorded using stitched images using the Nikon “Large Image Acquisition” tool, with 70% overlap between adjacent images in the vertical and horizontal directions of acquisition. After collecting the stitched images, the “Denoise.ai” feature of the NIS Elements software package was used to reduce the shot noise of the image data. The filtered stitched images were then exported as monocolor tiffs of each fluorescent image channel for data processing in MATLAB. For all experiments, FRET efficiency for each pixel of the image was calculated using Equation 1, where the I_D/D_ signal was the image data from Channel 1, the I_D/A_ signal was the image from Channel 2, and Equation 1 was applied pixel-by-pixel for the image. from Channel 2, and Equation 1 was applied pixel-by-pixel for the image.

### Microfluidic Origami Data Processing

#### 2-Inlet Device

2-Inlet Devices were imaged at the gradient chamber region of the device, as demonstrated in **Figure 3A**. 150mM and 200mM NaCl concentrations were used for both sensor concentrations flowed into the device in each of the two devices used (1 for each sensor type). Upon collecting the FRET response of the sensors at each gradient profile, a solution of red dye (McCormick) was then flowed in and fluorescence read using the Channel 2 laser settings. FRET response data was averaged at 6 regions of the device, corresponding to the 6 chamber inlet flows, over a region of 150 x 150 pixels (186 x 186 µm), with approximate size shown in Figure 3A. The red dye data was normalized from 0 to 1, averaged in the same spatial regions as the FRET data, and used to interpolate the NaCl concentration gradient utilizing Equation 3:

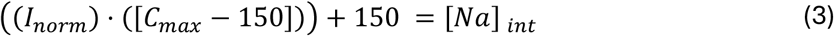

Where I_norm_ is the normalized dye intensity, C_max_ is the maximum concentration of the gradient (in this case 200mM), and [Na]_int_ is the resulting interpolated NaCl concentration for each point. Mean FRET values of each sensor type at each of the 6 points were then plotted versus the interpolated NaCl concentration and distance for each sensor type for comparison of spatial measurement, concentration sensitivity, and general applicability at large length scales.

#### 3-Inlet Device

3-Inlet Devices were imaged at the inlets of four different devices as outlined in Figure 4A. Each experiment was setup with three different NaCl concentration solutions flowing into the device, one at each inlet, with a sensor mixture at 150 mM NaCl used at the middle inlet of each experiment to ensure consistent concentration baseline. This resulted in four distinct concentration jumps for each sensor type of: Δ = 300mM, 200mM, 50mM, 25mM, for a total of eight groups. To assess sensor concentration and spatial resolution, 150 x 150 pixel (186 x 186 µm) regions were utilized for data processing at each fluid interface where concentration jumps occurred, as shown in Figure 4A. The 150 replicate traces across the region width (one for each row of pixels) were then averaged to generate a mean and standard deviation of sensor response. To assess both spatial and concentration resolution of each sensor type at each concentration profile, the maximum slope was calculated using MATLAB’s gradient function, which took the first derivate of the FRET response across space, with the max function finding the peak slope. This maximum slope was generated for all 150 traces, averaged for each group, and then compared across conditions. To determine the distance these changes in concentration could be measured over, the “detectable distance” was calculated for each replicate using Equation Y:

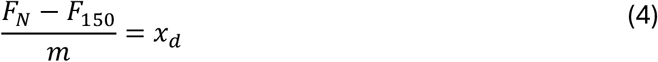

Where F_150_ is the mean FRET efficiency of the 150mM region, F_N_ is the mean FRET efficiency of the region of higher concentration, m is the maximum slope of the given trace, and x_d_ is the detectable distance. x_d_ provides a measure of the distance that these sensors can measure specific concentration changes over, with the regions F_150_ and F_N_ being defined as regions 10um away from where the gradient function was greater than 0.

## ACKNOWLEDGEMENTS

This work was supported by the National Institute of Arthritis and Musculoskeletal and Skin Diseases (R21AR076611 to BAW and CEC) and the National Science Foundation (CAREER 2143779 to BAW and 2323968 to CEC). We also acknowledge members of the Walter and Castro laboratories for useful feedback, and we acknowledge support from the OSU Campus Microscopy and Imaging Facility (CMIF) at The Ohio State University for TEM and Confocal imaging.

## CONFLICT OF INTERST

The authors declare no competing financial interests.

