## Supplemental Figures and Tables for "Tunable DNA Origami Nanosensors for Detection of Multiscale Spatial Ion Concentration Gradients"

**Supplemental Table 1:** Table of DNA sequences for the modulating and fluctuating linker for all four versions of the device. Helix 0, 3, 6, 9 and 12 are the modulating linkers and are therefore unconstrained when the corresponding unconstrained sequences are included or single stranded when the unconstrained sequences are excluded. Helix 15 is the fluctuating linker containing the donor-acceptor fluorophore pair with the controllable overhang length measured in base pairs.

| Name | Sequence | Description |
|---|---|---|
| 9bp-Cy3 | /5Cy3/ATTCTACTATCTGCGAACGAGTAGA TTTAGTTTGACCATTAGATACATT | Sequence containing the donor fluorophore for the 9bp overhang on the fluctuating linker |
| 9bp-Cy5 | TATATTTTCATTTGGGGCGCGAGCTGAAA AGGTGGCAT/3Cy5Sp/ | Sequence containing the acceptor fluorophore for the fluctuating linker for the 9bp version of the device. |
| 10bp-Cy3 | /5Cy3/AATTCTACTATCTGCGAACGAGTAG ATTTAGTTTGACCATTAGATACATT | Sequence containing the donor fluorophore for the 10bp overhang on the fluctuating linker |
| 10bp-Cy5 | TATATTTTCATTTGGGGCGCGAGCTGAAA AGGTGGCA/3Cy5Sp/ | Sequence containing the acceptor fluorophore for the fluctuating linker for the 10bp version of the device. |
| 11bp-Cy3 | /5Cy3/CAATTCTACTATCTGCGAACGAGTA GATTTAGTTTGACCATTAGATACATT | Sequence containing the donor fluorophore for the 11bp overhang on the fluctuating linker |
| 11bp-Cy5 | TATATTTTCATTTGGGGCGCGAGCTGAAA AGGTGGC/3Cy5Sp/ | Sequence containing the acceptor fluorophore for the fluctuating linker for the 11bp version of the device. |
| 12bp-Cy3 | /5Cy3/TCAATTCTACTATCTGCGAACGAGT AGATTTAGTTTGACCATTAGATACATT | Sequence containing the donor fluorophore for the 12bp overhang on the fluctuating linker |
| 12bp-Cy5 | TATATTTTCATTTGGGGCGCGAGCTGAAA AGGTGG/3Cy5Sp/ | Sequence containing the acceptor fluorophore for the fluctuating linker for the 12bp version of the device. |
| Unc. Helix 0 | TTTGAGGATTTAGAAGTATTAGACTTTACA AACAATTCGACAACTC<br>TCTAAAATATCTTTAGGAGCACTAACAACT AATAGATTAGAGCCGT | Top and bottom sequence for the unconstrained modulating linker on Helix 0 |
| Unc. Helix 3 | AGACGCTGAGAAGAGTCAATAGTGAATTT ATCAAAATCATAGGTCT<br>TTAACCTCCGGCTTAGGTTGGGTTATATA ACTATATGTAAATGCTG | Top and bottom sequence for the unconstrained modulating linker on Helix 3 |
| Unc. Helix 6 | TCGGCTGTCTTTCCTTATCATTCCAAGAA CGGGTATTAAACCAAGT<br>CCTGAACAAGAAAAATAATATCCCATCCT AATTTACGAGCATGTAG | Top and bottom sequence for the unconstrained modulating linker on Helix 6 |

| | | |
|---|---|---|
| Unc. Helix 9 | GTATGGGATTTTGCTAAACAACTTTCAACA GTTTCAGCGGAGTGAG ACTAAAGGAATTGCGAATAATAATTTTTTC ACGTTGAAAATCTCCA | Top and bottom sequence for the unconstrained modulating linker on Helix 9 |
| Unc. Helix 12 | CTTAGCCGGAACGAGGCGCAGACGGTCA ATCATAAGGGAACCGAAC ACGGAGATTTGTATCATCGCCTGATAAAT TGTGTCGAAATCCGCGA | Top and bottom sequence for the unconstrained modulating linker on Helix 12 |

**Supplemental Table 2:** Table of DNA sequences for all staples used to form the open and closed barrel of the NanoDyn structure when combined with the p8064 scaffold. The bottom barrel is the closed end while the top barrel is the open end.

| Sequence | Location |
|---|---|
| ATGCGTTCTAGCTGATAAACAGGAAATCCTGGAGGCCGCTAT | Cap End |
| TAATGAGAGATCTACAATCAATCAAAAGAGTAACTATC | Cap End |
| TTTCAGACAACACCAGTAATGGATCATCAGAGCTGGTCTGGTCAGTCCG | Cap End |
| CCTAAAGCGTTAAAAGGTTTTGACGTCACTGCTTGTAGAACGTCATTTC | Cap End |
| AGTGTTGGAACGCGTTAGTATCGCTCAAGAACGCGCCTGTTTATCAACA | Cap End |
| GAATAGCAATATATTAATTACTGAGAATACAACATGTTCAGCGGGTAAA | Cap End |
| AAAATCCTCTTCTGTAAGAATACGCCAACTGTCCAGACGACGTCATAAA | Cap End |
| GATGGTGGTTTGAAATACCCAGTCGGGACGAAAATTTTGGGGGCAA | Cap End |
| TGGAGCACTAGTGCCACGCCGGAGAGGGTAGGATAATCAATC | Cap End |
| GCGGTCCTAATGAAACTCACAACCGAGCGTTCTTCGCGTCCGCTGC | Cap End |
| TCAGAACTGAAAAACAGCAGGTCATTGCCTGGGTAATCAATT | Cap End |
| CTGATTGGGGCGCCAAGCATAGTGAAATTTTTCACGGTCATAGCGC | Cap End |
| AATCGCAAGACAAATTCCAGTTTGG | Cap End |
| TTTTTCACCGAGATAGGGCGATGGTCAGTAAAGATTCAAAAGGAAGCCT | Cap End |
| AATTTCACTTATAAGGGCGATATCAAACAAGGCCGGAGACAGCCCTGTA | Cap End |
| TTTAATGGTTCCGAACCATCATCACCTTACCATCAATATGATGTTGTAC | Cap End |
| TGTCTCACTGATCAGTTGAGGATCCCCGGGTTTAATTGGGCG | Cap End |
| TCGACCGTGTGATAAGAGGCACCGACAA | Cap End |
| GCCATGGTCATTTCTGCCAGCACGCCCCCTGTATTTACTCTGCTG | Cap End |
| TATAAAGCCAACATATGCGTTA | Cap End |
| GCCAGTACCCGCTTGGGAAGAAAGCCGGACGCCAGAGATTGT | Cap End |
| AAGGTAAGTGTGTTCAGCAAACGCAACCAGCTTACACCGATA | Cap End |
| CCTCAAAGAGAATACGGCATCCCGCCGCGCGCGTAATTAAAGGGCG | Cap End |
| GGGTAGCTGTTTCCTGTAAGTGTACCAGTAAAAGAATATCAC | Cap End |
| TCACTGTTGCCCTGCGGCCAATCCGCCGGGCGCGGT | Cap End |
| AATTAATGCACAGTAGGGCTTAATTAGAAAAAGCCTGTAGAAAAC | Cap End |
| GGTTTCTCATTGCAGGCGCTTTTATCAGTCGCTGAAACGAGGCATCTTT | Cap End |
| TCGACAATAACGCCATATTTAACAAAACACCGGAATCATTTAGTT | Cap End |
| CATCCCTGGTGTCCAGCATCAGAATTTCACGCATAAGAGGCTAAAAGAA | Cap End |
| CTGAGTAATTCATGTAATTTAGGCAATAAGGCGTTAAAACCTAAA | Cap End |

| | |
|---|---|
| AGCCGCTGCGACGAGCACGGGAGCCGCTATTGTCGGATTCTCCGTCGTT | Cap End |
| AGTGCTCAATAATATTAAGTAGAAGATTAAGAGCCAGC | Cap End |
| AACGCCGGTGCGTGCCTTCGAATTCGTAATCTAATGAGGCAG | Cap End |
| GGACGCGCCTCGGGCCGTGTTATC | Cap End |
| TCCAAAAGGAGCCTTAAAGGCCGCT | Cap End |
| GTTGCACAGGCGGCCATCCCATCGTTAATAAAGTATTTTCGA | Cap End |
| AGTCCGTAAAATTAAACGGGTACAATCGGCGGACATAATGCT | Cap End |
| ACGGTACATCAAACGTAATAACCTCACCGGAGCCTCTTTAAA | Cap End |
| GCACGCCAGCCCAGTCCGTGGAGCCGCCACGCTGGCGACGTT | Cap End |
| GTCCTTAAATGATAGACGCCAGTGCCAAGCTGCAAGGCGAACTCACTGT | Cap End |
| CCTCAGCAGCGATACCAAGCGCGAA | Cap End |
| GCATCGGGGCTTGCAGGGAGTTTAATTGTATCGGTTCGCACTTGGT | Cap End |
| CGGCTACACCGATATATTCGGCTTGCTTTCGAGGTGCGGGGTTTGC | Cap End |
| CTAAAGAAACAACCATCGCCCTTAAACAGCTTGATGGCTGGATACA | Cap End |
| AGTTTCCAAAAGCCGCGCCGACAATGACCTTTTTCCCAACCT | Cap End |
| GACCCCCGGCACCGCTTCTGGATCTGCCAGTTTGAATCATACAACG | Cap End |
| TACACTAAACCAGGCAAAGCGATGGGCGCATCGTAATTAGCATGCG | Cap End |
| AAAACGACCATTCAGGCTGCGGGATAGGTCACGTTAGCCTCAATTA | Cap End |
| GTTGTAAACGCCAGACATCACCGTTGTATGACCTGAAAACATGCAACAG | Cap End |
| CACTCCAGCCAACGACAGTATC | Cap End |
| ATCAACGGCGGATTGACATCAAATATTTAAA | Cap End |
| CCCACGCCAGGGAACGGCAGCGCCATGTTTAAGTTGGGCGTG | Cap End |
| TTAATGTGCTTTCAGAGCGGAATTTGTGAGATTCTGCTCCAG | Cap End |
| TTTCATCCAGCTCATAGCATGAGGCTATAGGTGAGGCGGTCAAGTC | Cap End |
| TGTTTGGGTAAACGACGTTTCTCCGTGGTGAAACA | Cap End |
| TCTGGCCGGAACGCGATGAACAGAGTCTCACCAGCAGAAGATTAGC | Cap End |
| AGTAGCATTAACATCCAATAAGGGGACGGCTTTCCAGCGATTAAAGACA | Cap End |
| TTGTAAAGGGAACAGGTGCGGAAAAATACGTAATGCCACTACTGGGAAG | Cap End |
| AATCCCCAAAAATTAATGCTGAGAGCCAGCATCGAGGTACGG | Cap End |
| CAAGTGTAGGTTGGCAAATCAACAGACTCCAACGTCAAAGGGTTG | Cap End |
| TTATTTCAGGCAAGGCAAAGAACCGTGCTGCCGGAAAACACTGTAGCAA | Cap End |
| GGAGGTGAGACCTCAATCAATATCAAAACCGTCTATCAATCAAAA | Cap End |
| ATACTTTAAATTAAGCAATAAGGTGTAGCCATTCGAAGAGGCTTGAGGA | Cap End |
| TGATCAAATCGCTGAACCTCAAATGGCCCACTACGTGAAATCGGC | Cap End |
| CAAAAACGAGCATAAAGCTAACGTAATGCAACTGTGAAGGCAATGAGGA | Cap End |
| ATAAGCGATTCAACAAAAATCTAAAGCACCCAAATCAAGTTTCCTGTTT | Cap End |
| ATGCAATGCCTGAGTAATGGATAAAAATTTTTAGAA | Cap End |
| TTTCACCGCCTGCAACAAATCGGAACCCTATGCAGCAA | Cap End |
| GGAAAGCGAAACCACACAGATGCCACCGTCGGTGGTGCCTTT | Cap End |
| CTACGATTTATTTTTATAGAAAAGATTTTGTTAAAATTACAA | Cap End |
| GGCAATGTGAGCTATCGGCCAACGCGCGGAGAGTTGGCTATTGTAT | Cap End |
| ATTTATTTTTCCGAGTATATGTACAATTTTTGTTAAATAACA | Cap End |
| ACCTACAGACATTCGAGCCGGAGGGTGG | Cap End |

| | |
|---|---|
| GGTAAAAATCGATGAAGGGTAAAGCGGCAGCGGGTTACTGAGCCT | Cap End |
| TTGCGTAACCAGGAGCGCGTTGCGCGTGCCAGCTGCATACGC | Cap End |
| CTTATGTGTAGCGGTCACAGCGGTCGCAAGAATGCCAATTAA | Cap End |
| GTAATATCATTTGCAACGCGGTCCGTTTGCGTGGTCGATCCACCGG | Cap End |
| AACCGTCTGACACACGAAAGCCTGGCGGTTTGCGTATTCCCT | Cap End |
| CAGAGAAGTGGAGCTTGGCCGTAATTTGCCCCAGCAGGAACC | Cap End |
| AGAGAGGCCAGAAAAGGGAGCCCCGGGCGCT | Cap End |
| GGCCTCTAAGGGGGAATGTGAGCGAGTACGCATTACCCGGTTCTAT | Cap End |
| AGTATCAGAGCGTATAACGGTTGTATGCTGATTGCCGTCAGC | Cap End |
| CCACGTGGCAAATGCGCCCGCCTGGCCCTGAGGGAGAGGGGT | Cap End |
| GCAGTAAAACTTTTTTAACCAATATTCC | Cap End |
| AACTTGCCTGCCGCCAGGATCAAATCGTCGCTGGCAGCTGCC | Cap End |
| TGAATAATTTGCACGTAAAACAAGCGCATTAG | Open End |
| TCACATCAATCAGTTCAGAAAACGCGAGAGGCTTTTGCAACCCTC | Open End |
| ACATAAATCAGAGAAATAGCAGAAACGCAACATATAAAAGAAATATGGT | Open End |
| GCAGCCTATTGAGTTAAGCACGATTTTTTGTTTACCCAATCCTAAGACT | Open End |
| CAAAAATTGAATATGATTATCCCATAAATCAAAAACGGATGGCTAAACC | Open End |
| AATTTTTACACGGAATTTTTACCCTGACTATTTTGATA | Open End |
| TTACCTTGTGAGTGTAAACAGAAGAACGATATAGAAGGCTTAATATTGA | Open End |
| CATTTAATGTAAATCCTAATTGAACCTCGCCCAATAGCAAGCTAAAGGT | Open End |
| TCAAGAAAATTTTCCTAACGAGTTTTGACATCGTAGGAATCAACCGACT | Open End |
| GCAAAAGAAGAAAAACATAGCGATA | Open End |
| GCTAATAAACAGGGAGAAATATTGGATTATCCCCCTCAAATGGTTTTGC | Open End |
| TTGCTTCCAATTTCAACGGATCAAAGAAAGAAGCAAAGCGGAAGAGTAC | Open End |
| ATTAATTAACAAAATGAATACTAACATTAAAAAGATTAAGAGACTCCAA | Open End |
| ATCCTTGTGATGAAGCGCAGATATTAATGAAAGACTTCAAATATCG | Open End |
| ACAATGAGATAACCCACAAGATTACAGAATTT | Open End |
| AATAAGAACCCAAAAAATACATACATAAGACAAAA | Open End |
| AATTTTATCCTATTAGTTGCTA | Open End |
| AGTTACCAGAAGGAAACCGAGATAGCTATCTTACCATTGAGC | Open End |
| CCTTACGGTATTCTCCATATTATTTATCATAAATCAATATATTTTT | Open End |
| TTAGCAACGATTGAGGGAGGGCAGACTGTAGCGCGCATTTCGTCACCA | Open End |
| TCGAGAACAAGCAAGCCGTTTATTAGAGCCAGCAAAATC | Open End |
| TCAATAGAAAATTCACGCAAAGACACCA | Open End |
| TTACCAGCCTTATTAGCGTTTCTCCCTCCGCCGCCACAAATAGTAAGCG | Open End |
| AAAAGGTGGCAATAATAACGGAATGCAAGAA | Open End |
| GGGCGACCGGCATTTTCGGTCCAGAACCCACCAGACCAGAAT | Open End |
| CGGAAATAATCAAGTTTGCCTTACAACGAGGAACCTTGCTCAACCGGAT | Open End |
| CATAAATCAGCGAGGCGTTTTAGCTGCCAGTTACAAAAAATAACC | Open End |
| GAATTATCACCGTAATCAGTACAGACAGTTCAGGGAGTGCCGTCAAGAG | Open End |
| GTCTTACCGCCCGACTTGCGGGAGGCGTCTTTCCAGAGCGTCGCT | Open End |
| TGAGCCACAATGAAACCATCGGTAACGACTCAGAGATAAGTAGCATAGG | Open End |
| GGATTATTTTAGCCTTAAATCAAGGAATCTTACCAACGCCTTAGA | Open End |

| | |
|---|---|
| CATATTCAACACGTAGAAGAACTGGCATGATAAATAAGAACCAAT | Open End |
| CCATTAGCAAGGCCGGAAGTCTTTCGCCACCCTGTATCATGAACGG | Open End |
| TACAAACTTAGCGTAAGGTAATCTTACGCAGTATG | Open End |
| CTGAGCCAGGATTATAAGAGGCTGAGACCCTATTTCGGAACCATAAAAC | Open End |
| CCCTCAGAACCCAGACGTTAGTAAA | Open End |
| CTTGATATTCACAAAGCATTGACAGGAGACCACCG | Open End |
| ATTAAAGACCACCACCAGAGCAGAGCCGCCACCCTATAGCCCCGCC | Open End |
| GGAAACTCAAGAGAACCCTCAGAGCCGCGCCACCCTCAGAGCTTTT | Open End |
| GCGGATAATAGCAAGCCCAATCCTGTAGCATTCCAGCGACAGTATT | Open End |
| GGTTGATCCACCACCCTCATTCCCTCATAGTTAGCATAGCAGCACC | Open End |
| GAATAGGTCAGAACCGCCACCTCTAAAGTTTTGTCACGTCACTTTG | Open End |
| TCATACAGCCTTGAGTAACAGTTCATCAGTTGAGATTTA | Open End |
| AATTTACATAAACAGTTAATGACATTATTACAGGTCTATCATAAAA | Open End |
| ATACATGAAAGTATGGATTAGCGGGGTTCATGTACCGTAACA | Open End |
| TAATCTTCGTAACAAAGCTGCTCATTATACCAGTCGCTCAACTGCT | Open End |
| CTGGCTGGTGAATAAGGCTTGTATGCGATTTTAAGCAACTAAGGAT | Open End |
| TGTACAGGAGAAACACCAGAATTTAATCATTGTGAAAGTTTCGAAG | Open End |
| TGAAAGAGGACAGACCGTACTCAGG | Open End |
| CTGAATCATTACCCTGGGAAGAAAAATCATTGCTGATTTTTGTCAG | Open End |
| GAACTAAAGACGACGATAATAGAGCTTATACGTTATATTATT | Open End |
| TTAATTTCAACCGAGTAGTAAA | Open End |
| GGAATTACGAGGCAGGTAATATCATTGAATACTTC | Open End |
| GTTTACCCGGAACACCCCCTGTCGCGCAGTCTCTG | Open End |
| CAGAGGGTAGTAAGAGCAACAAGAAAGATGCCCGTCGTTCCAAATCCTC | Open End |
| GAACTTTAAAATAATCCTGATTGTAAGAAATTGCGTAGGAGAATA | Open End |
| AAAATAGAGAATGAAGATGATGGCAATTGGTTTAACGTCAGAGAAAATA | Open End |
| AGAGGTCAATATAATGCTGTAAGGACGTAAATCAAGACAAGAGTACCAG | Open End |
| CCTTATAGTCACCACCAGAAGGAGTCGGGAGAAACAATATTTGAA | Open End |
| CTTTAATATGTTTTAAATATGAACTGGCTCATTCAACCTTCATCGAGAG | Open End |
| TAGTTGCATCATCATTTTGCGGAATCGCCTGATTGCTTTTAATTA | Open End |
| CAGGTCAAGTACGGTGTCTGGATTACCTCCCTGACACCAGGCTAGCCC | Open End |
| CAAGAAGCCCTTTAAAAGTTTGAGCAAGTTACAAAATCACAAACA | Open End |
| TCAAAGCGAACCAGACCGATTCCATATAACAGTTGATTC | Open End |
| GTCATCATCATATTCCTACAGTAACAGTACCGGAAACAGTAACGT | Open End |

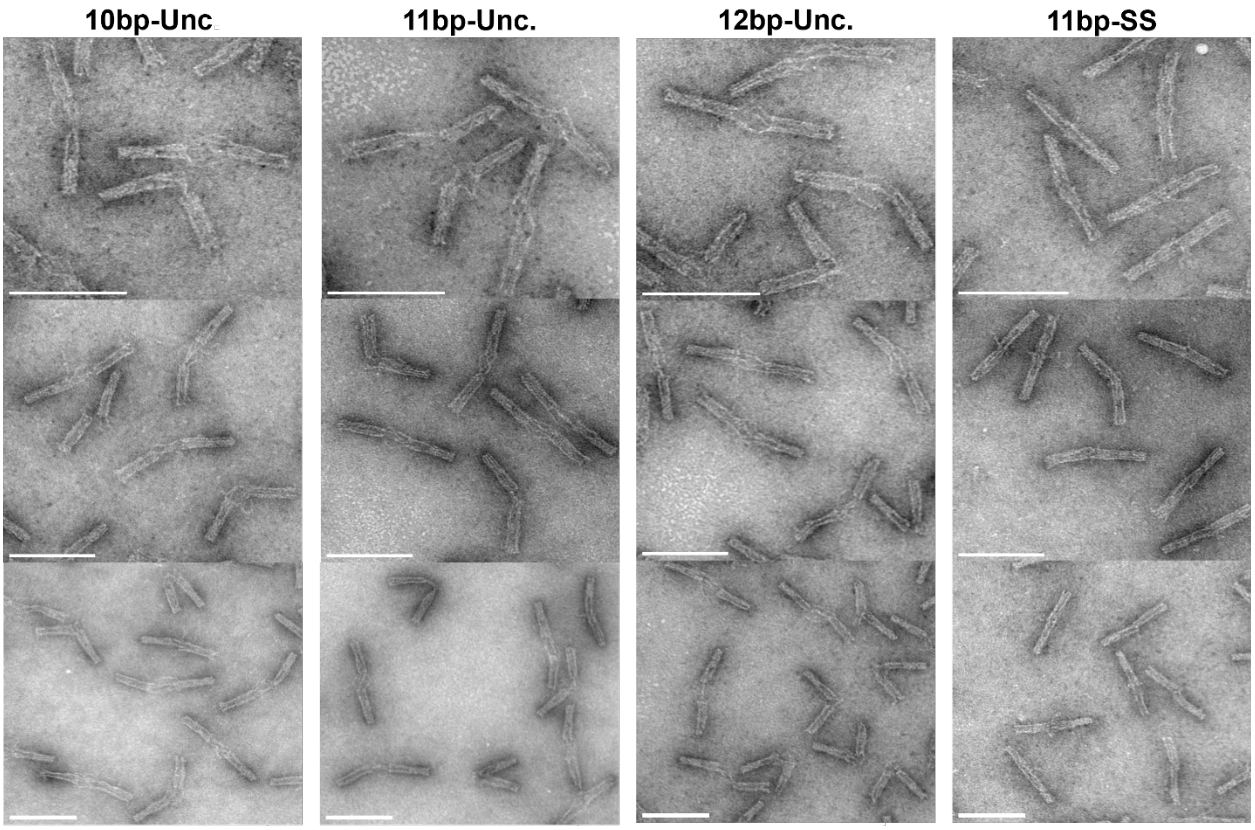


**Supplemental Figure 1:** Images of the different versions of the DNA origami structures tested at varying magnifications to visualize the stable state of the structure. The magnification that each image was taken at starting from top to bottom is: 150kx, 100kx, and 60kx. As explained in the text and shown in the image, as more base pairs are added going from left to right, the structures tend to favor the closed stable state over the open stable state. Each scale bar represents 100 nm.

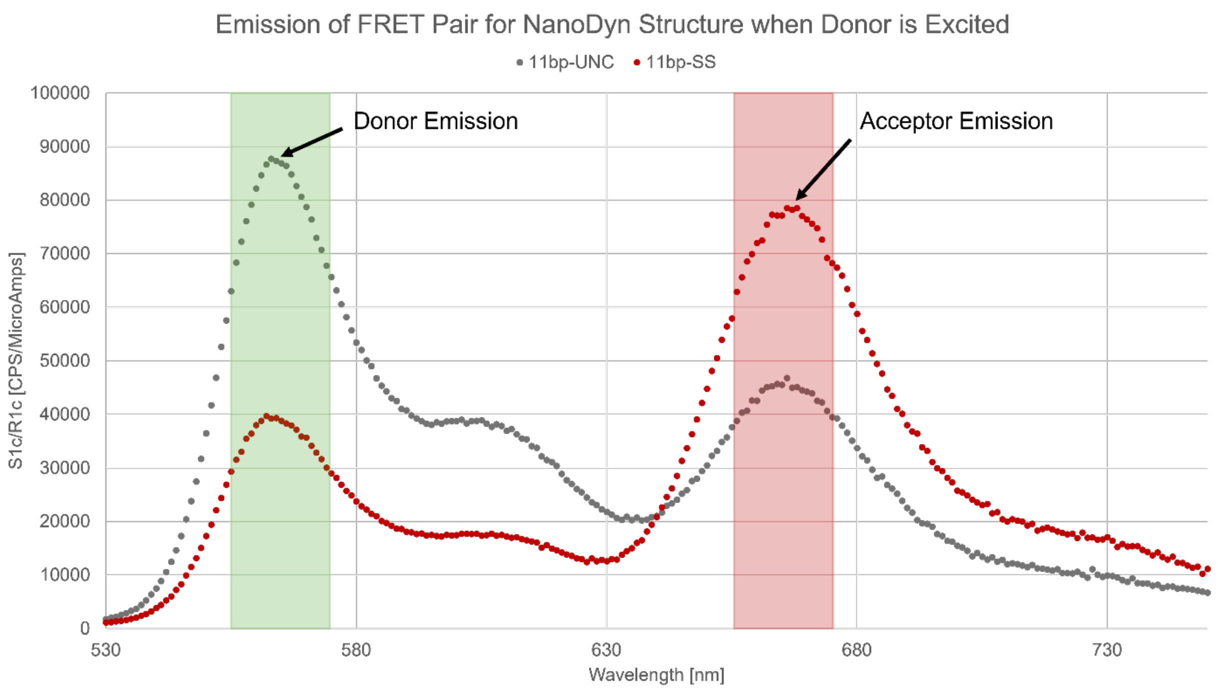


**Supplemental Figure 2:** Graph of the FRET pair emission on the fluorometer when the donor is excited directly at 510 nm. The graph shows the difference in emission peaks for the two variations of the device: 11bp-UNC and 11bp-SS. As the device shifts from the open configuration to the closed configuration, the acceptor emission peak overcomes the donor emission peak. Therefore, the 11bp-SS device favors the closed state more than the 11bp-UNC version which is confirmed by our TEM images.

**Supplemental Table 3:** Sensor Design Parameters Effect Sensitivity to Salt Concentration

| Design Version | Initial Sensitivity (dFRET/dC @ 100mM) [$1x10^{-3}$/mM] | Saturation Constant ($C_{sat}$) [mM] | Effective Range (Max: 100+2*$C_{sat}$) [mM] |
|---|---|---|---|
| 10bp-UNC | 0.19 ± 0.03 | 788 ± 133 | 100 – 1675 |
| 11bp-UNC | 1.23 ± 0.24 | 133 ± 9.6 | 100 – 365 |
| 12bp-UNC | 1.45 ± 0.12 | 174 ± 16 | 100 – 448 |
| 11bp-SS | 3.13 ± 0.50 | 69 ± 6.6 | 100 – 238 |

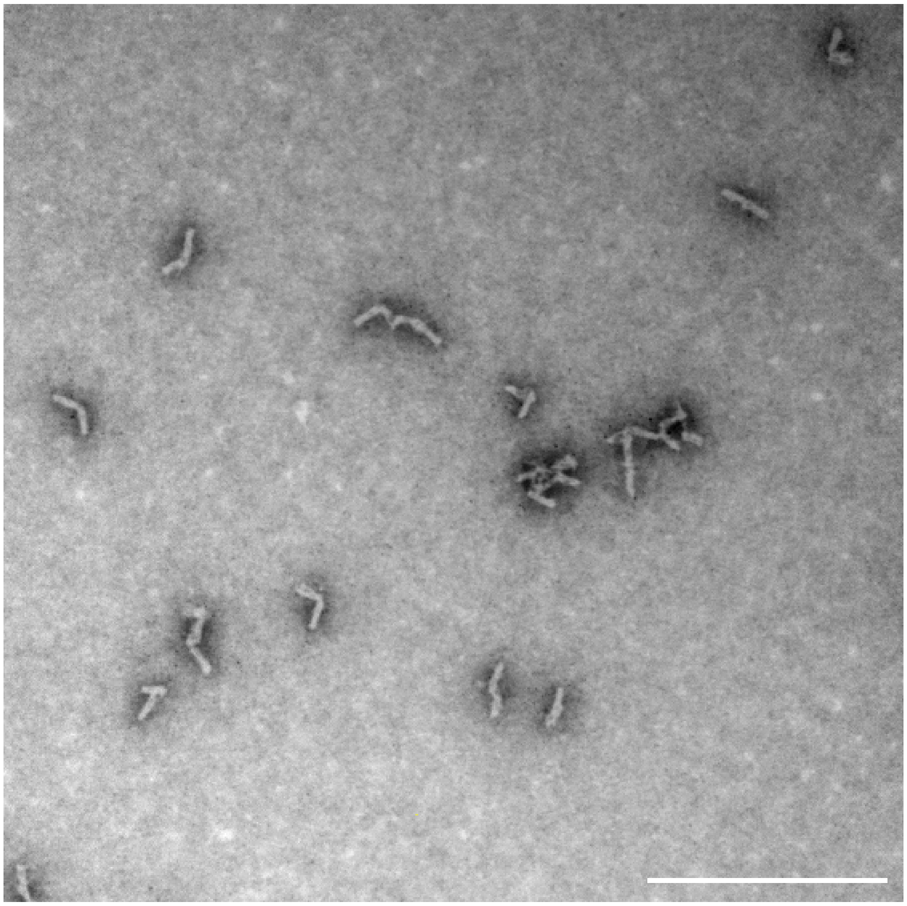

**Supplemental Figure 4:** Image of 11bp-UNC DNA origami structures from the outlet channel after being flown through the microfluidic device with inlet conditions of 450-150-150 mM NaCl + 1mM $MgCl_2$. This image confirms that the devices remain intact throughout the duration of the experiment. The scale bar is 500 nm.

**Supplemental Table 4:** Sensor Design Parameters Effect Spatial Sensitivity to Salt Concentration

| Concentration Profile (mM) | Concentration Change (mM) | Max Slope (FRET Eff*$10^{-3}$/μm) | | Distance to Equilibrium (μm) | |
|---|---|---|---|---|---|
| | | 11 UNC | 11 SS | 11 UNC | 11 SS |
| 150 to 450mM | Δ300mM | 11.39 ± 4.09 | 3.36 ± 1.0 | 6.56 ± 1.86 | 12.66 ± 3.06 |
| 150 to 350mM | Δ200mM | 6.90 ± 2.53 | 4.36 ± 1.50 | 7.69 ± 2.72 | 9.15 ± 3.29 |
| 150 to 200mM | Δ50mM | 2.00 ± 0.51 | 2.57 ± 0.73 | 9.32 ± 1.84 | 9.93 ± 2.88 |
| 150 to 175mM | Δ25mM | 1.23 ± 0.39 | 1.56 ± 0.33 | 7.67± 2.53 | 9.51 ± 2.75 |
